# Using a diabetes discussion forum and Wikipedia to detect the alignment of public interests and the research literature

**DOI:** 10.1101/496927

**Authors:** Fereshteh Didegah, Zahra Ghaseminik, Juan Pablo Alperin

## Abstract

**Background:** Diabetes is a chronic disease that affects millions of people worldwide. It is therefore unsurprising that there is a high volume of public discussions, resources, and research tackling various aspects of the disease. Over the last decade, more than hundred thousand research articles have been published by researchers and countless of online discussions have taken place on various online platforms. This study is an attempt to identify the areas of public interest, related to diabetes, by looking at online discussion forums and to evaluate their relationship to pages about diabetes found on Wikipedia and to the academic research about the topic. The main aim is to investigate the extent to which researchers are responding to the public’s interests and concerns, and to the level of uptake of the research topics in the public sphere.

**Methodology/Principal findings:** To detect public interests and concerns in diabetes, we collected posts on a popular diabetes discussion forum (DiabeticConnect) and pages (articles) about diabetes published in Wikipedia. We also downloaded the titles and abstracts of research articles about diabetes from the Scopus database, all between 2008 and 2016. Tags assigned to each post in the discussion forum were used along with the post itself to compute a Labeled Latent Dirichlet Allocation (LLDA) model, which was then used to classify the Wikipedia pages and research articles. The resulting classifications were then used to compare the prevalence of the topics found in the discussion forum with those of the other two sources. The results show that while research articles and Wikipedia pages about diabetes focus on diabetes testing, treatments, and disease control, the public forum discussions focus on Type 2 diabetes, emotional support, and proper diet for diabetic patients. However, for some other topics there was an alignment in the relative rise and fall of interest across the three platforms.

**Conclusions/Significance:** The alignment and misalignment in the changes of relative interest over the various topics is evidence that the LLDA modelling can be useful for comparing a public corpus, like a diabetes forum, and an academic one, like research titles and abstracts. The success of using LLDA to classify research articles based on the tags assigned to posts in a public discussion forum shows that this a promising method for better understanding how the scientific community responds to public interests and needs, and, on the flip side, how the public takes up the language and topics discussed by the academic community.

## Introduction

Diabetes is a serious health problem that has nearly doubled in prevalence among adults in the last three decades. All told, in 2012, 3.7 million people died from diabetes and high blood glucose (WHO, 2012). As the prevalence of the disease grows, so has the wealth of information that is available about it online; although some clinicians and health care professionals warn about risks of misinformation on the web and online platforms (Murray et al., 2003), patients are more inclined to search online than to ask their doctors when looking for support and guidance in making health care decisions (Greene et al., 2011).

Of course, not all online sources are the same. Some spaces, like online community forums, have been found to be largely free of misinformation (Balkhi et al., 2014). Such forums have become spaces where diabetes patients share their experiences, seek information, ask for help, and receive support from others who have the same health concerns (Hilliard et al., 2015; Greene et al., 2011; Ravert, Hancock, & Ingersoll, 2004). Moreover, research shows that online social ties, such as those that develop when interacting in an online forum, can have a positive impact on improving health and decreasing health-related anxiety among patients with chronic diseases, including diabetes (Hilliard et al., 2015; Balkhi et al., 2014; Sarasohn-Kahn, 2008).

As patients and the general public access information and find value in online discussion forums, we posit that their posts and discussions can also be valuable sources of information to learn what the public is interested in and concerned about. This information may in turn inform a broad community including patients, practitioners, researchers, civil society organizations, and private firms in deciding policies and taking appropriate actions to improve health outcomes for those affected by diabetes. As such, the present research aims to use online discussions around diabetes to identify areas of public interest or concern and to compare them to those being discussed by researchers in scholarly publications. Doing so will shed light on the extent to which researchers are responding to the public’s interests and concerns, and to the level of uptake of the research in the public sphere.

## Literature review

### Research alignment with public needs

This work is not the first attempt to use online communities to understand public interests in health issues. Tools like ‘*Healthmap.org*’ and ‘*Google Flu Trends*’ collect health-related information from different online sources such as news outlets, government newsletters, and web searchers to monitor public health problems (Brownstein & Freifeld, 2007). Google has recently started a project, ‘*Searching for health*’, to develop a tool^1^ showing how Google searches for health information correlate with actual incidences of diseases. The tool covers information about several highly searched diseases such as obesity and diabetes. The correlations show a rather strong relationship between Google searches in an area in the US and occurrences of diabetes in the same area. However, such online information has not been used to measure the alignment of public interests with the work being done by researchers, nor the relationship between both groups. Poor interactions and communication between science community and rest of society may cause scientists to remain uninformed of public demands in specific areas. Hence, public engagement in research projects is widely encouraged, especially for health-related topics (Marris & Rose, 2010), including an increased emphasis on involving the public in the research process even before projects start (Boëte, 2011).

For public health concerns like obesity, the alignment/misalignment between the research community and societal interests has been explored. Cassi et al. (2017) explored societal demands through the questions received by the European Parliament between 2009-2014 and found that a few topics identified by the public, such as sugar and food economy, are ignored by the scientific community and also that obesity research mainly focuses on biomedical science while largely neglecting socio-economic factors (Cassi et al., 2017). Similarly, misalignments were found between public health issues measured through disease burden rate and the amount of research done on the same issues. For example, depression or stroke were found to be prevalent issues around the world but there is minimal research done in these areas (Rafols & Yegros, 2018). Also, no correlation was found between the rate of global disease and disability burden and number of medical articles published in MEDLINE that are relevant to those diseases. Moreover, the health research publications were not relevant to the health issues in poor countries (Evans, Shim, & Ioannidis, 2014).

Another study explored whether pharmaceutical research published by the top 23 pharma companies was associated with a reduction in disease burden rates and found a mismatch between burden of diseases and the amount of relevant research published by the pharma companies (Yegros, et al., 2018). Moreover, Yegros et al. (2018) found no alignment between pharmaceutical research (with a disproportionate focus on cancerous tumors) and global health trends (where infectious and parasitic diseases are the main causes of deaths worldwide).

### Forums and Wikipedia

While comparing research to disease prevalence provides a window into topics of public interest, other sources are needed to capture the public’s concerns over time. This study suggests online discussion forums and Wikipedia as valuable sources that can provide up-to-date information on what the public is concerned about. In the case of diabetes, the increasing number of discussion forums focused on the disease clearly shows the public’s interest in discussing their concerns in such online platforms (Hilliard et al., 2015). Similarly, popularity of Wikipedia, currently the 5^th^ most accessed page on the Web (Alexa, 2018), makes it an important place to observe what topics the public consults and contributes when seeking information about a wide range of topics, including health, in general and diabetes, in particular. We are not aware of previous research that examined these platforms as sources of information about public health concerns.

Unlike many social media platforms, forums functionality allows high levels of engagement and interactions. Forums are usually focused on narrow topics, such as specific health problems and diseases and participants are mainly those affected, either directly or indirectly, by the concern. On health-related forums, members can ask questions and receive answers from other members who range from patients or family members of those affected to health practitioners and experts who contribute to the discussion. Patients also often use forums to discuss complementary or alternative medicines that their physicians may be less aware of (Eysenbach, 2003).

Online communications through different platforms, including forums, have been debated by physicians due to increasing rate of misinformation found online (Ahmad et al., 2006) and the potentially hurtful communication that takes place there (Crocco, Villasis-Keever, & Jadad, 2002). However, Balkhi et al. (2014) found no signs of misinformation on diabetes forums. More than half of parents who had a child suffering from diabetes signed up on the forums to receive/share information and they believed their participation in the forums helped them to better take care of their children (Balkhi et al., 2014). Some parents used forums for gaining specialized knowledge from experts and researchers (Greene et al., 2011) and patients found forums the most suitable place to express their feelings (Ebrahim, 2009).

In a study of forums for youth with diabetes, around 49% of the posts contained requests for information and another 50% of the posts sought social support, indicating the role of forums as more than sources of information. Other topics such as medical care, disease management, psychological effects, and, naturally, factual information about diabetes were also discussed in the forums (Ravert, Hancock, & Ingersoll, 2004).

Rolia et al. (2013) suggest that medical forums are also important sources of information for medical experts and healthcare providers, as they provide the opportunity to stay informed of health issues and concerns from a patient’s perspective, which in turns allow them to consider patients’ perspectives in their practice. This type of use has been especially effective in forums with the ability to allow patients registered in a personal health record portal to access related forums, find most relevant groups and topics, and browse relevant information (Rolia et al., 2013). While health practitioners in Rolia et al.’s study would personally read forums to understand patient perspectives, our study explores automated ways of gathering the collective concerns of forum members.

In a very different way than forums, Wikipedia is another popular source for health information and is widely used by the general public, health professionals and researchers alike (Herbert et al., 2015). It is the common starting point for patients to look for their required health information (Thomas, Eng, de Wolff, & Grover, 2013), and for good reason: in searches for medical research articles, Wikipedia was ranked highest in Google in searches for both general health and rare diseases (Laurent & Vickers, 2009). Although the reliability of Wikipedia has been debated and people are asked to critically read Wikipedia pages about health related topics (Hasty et al., 2014), Viseur (2014) found the platform comparable to its commercial competitors and also comparable to peer-reviewed platforms for the effectiveness of the peer-production model that allows readers and a community of editors to detect and correct errors quickly. As a result of its wide adoption and its model, Wikipedia plays a synthesizing role, where the public has worked collaboratively to summarize what is known about different aspects of diseases like diabetes. Its use as a source of health information is likely to continue, as clinicians continue to recommend patients contribute to Wikipedia instead of spreading their time and energy across a wider set of tools (Heilman et al., 2011).

### Research questions

Taken together, online discussion forums and Wikipedia capture a significant amount of user-generated content about health topics like diabetes. This study therefore seeks to examine the nature of this content for the purpose of understanding the topics that are of greatest interest to the public, and their relationship to the topics most explored by the scientific community. By analyzing the relationship between a diabetes forum, Wikipedia, and research articles, this paper aims to not only uncover the alignment or misalignment in topics, but also to understand who is driving discussions about topics: the public, or academics? In particular, this study investigates to what extent discussions on the public online forum DiabeticConnect and on Wikipedia align with academic research in diabetes found in the Scopus database. In doing so, it answers the following related questions:

1. To what extent do the topics discussed on the online forum DiabeticConnect align with the diabetes topics most edited on Wikipedia and with research papers about diabetes found in the Scopus database?
2. Are the topics and language of the online forums later found in the research in ways that suggest researchers are tuned in to public interests? Or is it the other way around?

By answering these questions through the use of topic modeling techniques, this work points to a new class of indicators that have the potential to create a greater understanding of how research circulates and influences the public sphere, so that research agenda and knowledge mobilization strategies can be changed in ways that leads to better health outcomes.

## Methods

Three types of data, forum discussions, Wikipedia edits and research articles, were derived from three sources: the diabetes forum *DiabeticConnect^2^*, Wikipedia, and the Scopus database. Data collection from all sources was carried out between July and August 2017.

### Forum

DiabeticConnect is a diabetes discussion forum with free membership for everyone with any questions or concerns about diabetes. The forum helps patients connect to other patients and experts and researchers in diabetes and at the time of data extraction had collectively 26,845 posts between 2008 and 2016. Each discussion thread is assigned at least one ‘tag’ by forum moderators to reflect the topics of the discussion. Because these tags are assigned by moderators and not by the members themselves, they are applied across threads in a relatively consistent manner and provide the basis for understanding the topics discussed in the forum.

With the forum moderators’ permission, we extracted all the discussion posts from the forum that included the text of the post, the date it was made, and the tags assigned. We restricted these posts to the 26,845 that were published between 2008 and 2016. We manually scanned the list of tags that were assigned to each post to normalize obvious variants such as ‘type 2 diabetes’ which could also be found as ‘type 2’ and ‘type ii’. We followed this procedure for the 100 most frequent tags to ensure that every instance of each topic was counted when we subsequently selected 10 most frequently used (Table 1).

**Table 1.** Most common tags on DiabeticConnect forum.

| **Tag** | **No. of**<br>**discussions** | **%discussions** |
| --- | --- | --- |
| Type 2 | 7178 | 26.74 |
| Emotional support | 3165 | 11.79 |
| Diet | 2808 | 10.46 |
| Type 1 | 2585 | 9.63 |
| Diagnosis | 1627 | 6.06 |
| Treatment | 1447 | 5.39 |
| Weight loss/Obesity | 1348 | 5.02 |
| Medication | 1291 | 4.81 |
| Control | 1071 | 3.99 |
| Blood sugar | 808 | 3.01 |

### Wikipedia

Wikipedia is an online encyclopedia that allows readers to collaboratively create and edit pages on any notable topic. By searching for the word stem “diabe*” in the Wikipedia API^3^, we identified 693 articles on the English edition of Wikipedia. We then filtered out 484 articles that were about celebrities who had died from diabetes using the list found in the Wikipedia page “Deaths from Diabetes” (Wikipedia, July 2017). We subsequently queried the remaining 207 articles in the Wikipedia API for the content of the page along with their edit history, including the number of deletions, additions and total number of edits made to the page. In total, the 207 pages were edited 23,627 times, with an average of 17.42 edits per page, per year (median: 8; sd: 25.07), and 16.95 edits per page over the 9 years studied (median: 8; sd: 24.64).

### Scopus

Scopus is the largest abstract indexing database in the world with over 69 million records from more than 36,377 journals. We identified 108,180 articles from Scopus by searching the online portal for articles with the word stem ‘diabe*’ in their title, abstract and author-supplied keywords with a published date between 2008 and 2016. The title, abstract, and published date was retrieved for each article. Titles were on average 13.27 words (median: 13; sd: 5.39) and abstracts on average 203.43 words (median: 205; sd: 74.96).

### Topic Modelling Procedure

Because our primary interest was to understand the public’s concerns with diabetes, we used the forum discussions as the basis for understanding the topics that the public is interested in. Using these tags as labels and the text of the discussion posts as the content, we computed a Labeled Latent Dirichlet Allocation (LLDA) model (Ramage, Hall, Nallapati, & Manning, 2009). Topic models are a type of text-mining tool that uses word frequencies and co-occurrences (when two words are found in the same document) to produce clusters of words that have a high probability of being found together within given corpora (i.e., set of documents). Unlike their unlabeled counterparts (Latent Dirichlet Allocation, LDA) that begin with a set of words and produces an unspecified number of word clusters (without labels), LLDA models begin by grouping documents into a fixed set of clusters (one for each label) and then produces a set of words most likely to be associated with each. In both types of models, each word is also assigned a weight relative to that word’s contribution to the cluster. While an LDA model would be appropriate for uncovering latent topics found in the discussion posts, an LLDA model is more appropriate for uncovering which are the words most likely to be associated with each of the tags assigned by the moderators, as well as the relative weight of each word.

We thus used the 10 most frequent tags found in the discussion forum (Table 1) and computed the LLDA model (i.e., the list of words and associated probabilities) using Mallet software^4^. This model was then used to classify each of the documents in the other two corpora (i.e., Wikipedia and Scopus). As a result, every Wikipedia page and research article were assigned the tag from the forum whose words (as per the computed LLDA model) most closely resembled their content.

### Lead/lag visualization

To observe the interest in each of the topics, we plotted the number of posts, edits and articles with each of the tags as a percentage of the total number of forum posts and research articles, and as a percentage of the total number of Wikipedia edits. It was necessary to use number of Wikipedia edits as a measure of interest in a topic given the dynamic nature of Wikipedia, and the relatively rarity of creation of new Wikipedia entries related to Diabetes. These values were calculated for every year, using publication date for the forum posts and the research articles, and the date of the edit on Wikipedia.

These time series are used to depict the relative rise and fall in activity of each topic across the three platforms. We were interested in seeing if changes in the relative interest of one topic led to a change in the others (e.g., if the public’ growing interest in a topic led to more research about it, or vice versa). To better compare such lead/lag patterns between the three platforms, we normalized each time series by subtracting frequencies of the topic over years by the mean frequency and then dividing by the standard deviation. These normalized graphs were then manually shifted back and forth in time to find the best alignment, similar to the strategy used by Shi et al. (2010).

## Results

### The alignment between the forum, Wikipedia and research topics

Health is the most frequent topic extracted from both research articles (around 25% of articles are classified under health) and Wikipedia articles (around 30% of articles are classified under health), but since it is a very general topic and does not specifically relate to diabetes, we excluded the topic from further analysis. With that omission, the top five forum tags assigned to research articles by our LLDA model were, in descending order of frequency: A1C test, diabetes treatment, diabetes control, type 1 diabetes, and diabetes complications. The top five tags assigned to Wikipedia pages were, in descending order of frequency: diabetes control, type 1 diabetes, diabetes treatment, A1C test, and insulin. And on the DiabeticConnect forum, the five most frequent tags were: type 2 diabetes, emotional support, diabetes diet, type 1 diabetes, and diabetes diagnosis (Table 2).

**Table 2.** Top 10 topics across the three platforms.

| Forum |  | Wikipedia |  | Research |  |
| --- | --- | --- | --- | --- | --- |
| Topic | % discussions | Topic | % edits | Topic | % articles |
| Type 2 | 26.74 | Control | 11.97 | A1C test | 16.81 |
| Emotional support | 11.79 | Type 1 | 11.12 | Treatments | 13.47 |
| Diet | 10.46 | Treatments | 10.13 | Control | 13.16 |
| Type 1 | 9.63 | A1C test | 8.86 | Type 1 | 9.60 |
| Diagnosis | 6.06 | Insulin | 4.61 | Complications | 6.05 |
| Treatments | 5.39 | Blood sugar | 4.13 | Awareness | 3.09 |
| Weight loss/Obesity | 5.02 | Symptoms | 4.04 | Symptoms | 2.93 |
| Medication | 4.81 | Complications | 3.95 | Neuropathy | 1.84 |
| Control | 3.99 | Diet | 3.73 | Insulin | 1.71 |
| Blood sugar | 3.01 | Awareness | 2.99 | Diet | 1.21 |

The topics on each platform already show that not all the highly discussed topics on the forum are of interest to researchers, nor to the people who edit Wikipedia. While a high percentage of research articles and Wikipedia are about diabetes testing (especially A1C test), control and treatments, a large number of forum posts discussed emotional support and motivation for patients and diabetes diet, topics which are not found among the most popular topics in the research literature. On the other hand, the most edited topics on Wikipedia have greater overlap with the topics of research articles.

### The lead/lag between public discussions, research and Wikipedia

We traced the prevalence of some of the most discussed topics in the forum over time and compared it with the prevalence of those topics in the research found in Scopus and on the Wikipedia pages about diabetes. We plotted the relative interest in the topics over time across the three platforms for the top ten most discussed forum topics. These time series allowed us to visualize whether the rise and fall in the interest of topics in the discussion forum lead or lag similar changes in the research literature and in Wikipedia articles about diabetes.

We normalized the values for each year by dividing the percentage of posts and articles by the average across all years and plotted both the standard percentages (Figures 1a-j, left) and the normalized values (Figures 1a-j, right). These figures show that in six of the 10 topics (‘emotional support’, ‘diet’, ‘type 1 diabetes’, ‘diagnosis’, ‘medication’, and ‘blood sugar’), the time series align across the three platforms, with corresponding peaks and valleys for each, indicating that changes in public interest on these topics are accompanied by similar changes in what research is published and what topics are edited on Wikipedia.

We found a lead/lag pattern for two of the topics (‘type 2 diabetes’ and ‘diabetes treatments’) (Figures 1a & 1j). In the case of ‘type 2 diabetes’, the normalized and shifted time series show that Wikipedia edits lag forum discussions by one year while research articles lead forum discussions by a year. This shows that forum users had increased interest in type 2 diabetes one year ahead of editors of Wikipedia pages, but that researchers had increased interest in the topic one year ahead of forum users. In the case of ‘diabetes treatments’, forum discussions lead the research by one year and the Wikipedia edits by two years. That is, forum users had increased interest in discussing treatments for diabetes in 2009, and, one year later in 2010 there was a corresponding rise in the number of articles that took up the language found in the forum. Then, the following year, the Wikipedia pages that used similar language were edited more heavily (peaking in 2011).

The remaining two topics had a different pattern. Posts and articles about ‘weight loss/obesity’ show alignments across the three platforms between 2008 and 2011, but diverge in the following years (Figure 1f). From 2012 onwards, forum discussions seem to lag both research published and Wikipedia edits. Wikipedia edits about articles related to ‘weight loss/obesity’ dramatically increased in 2012 when academic research and forum discussions had their lowest number. The pattern is the reverse for ‘diabetes control’, where the three platforms do not align prior to 2012, but all show increased interest in the topic starting in 2012 (Figure 1h). For this topic, forum posts and research articles are relatively aligned with low levels of activity, while Wikipedia edits on pages associated with the topic show a different pattern, rising dramatically in 2010.

**Fig 1:**
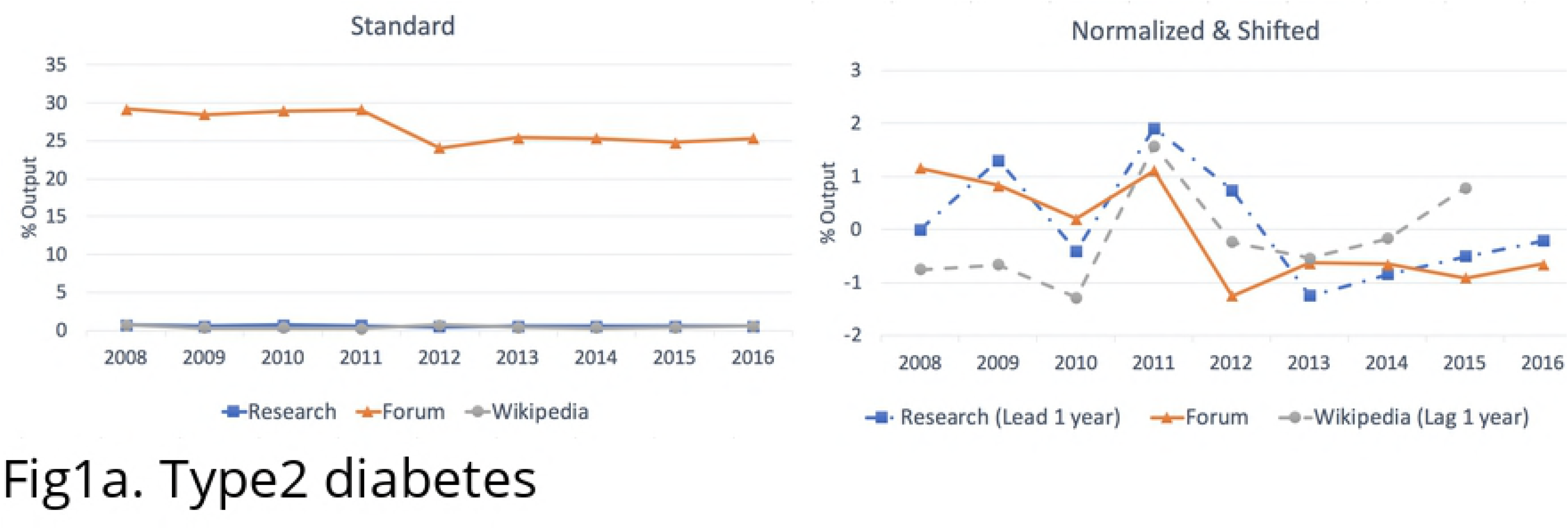
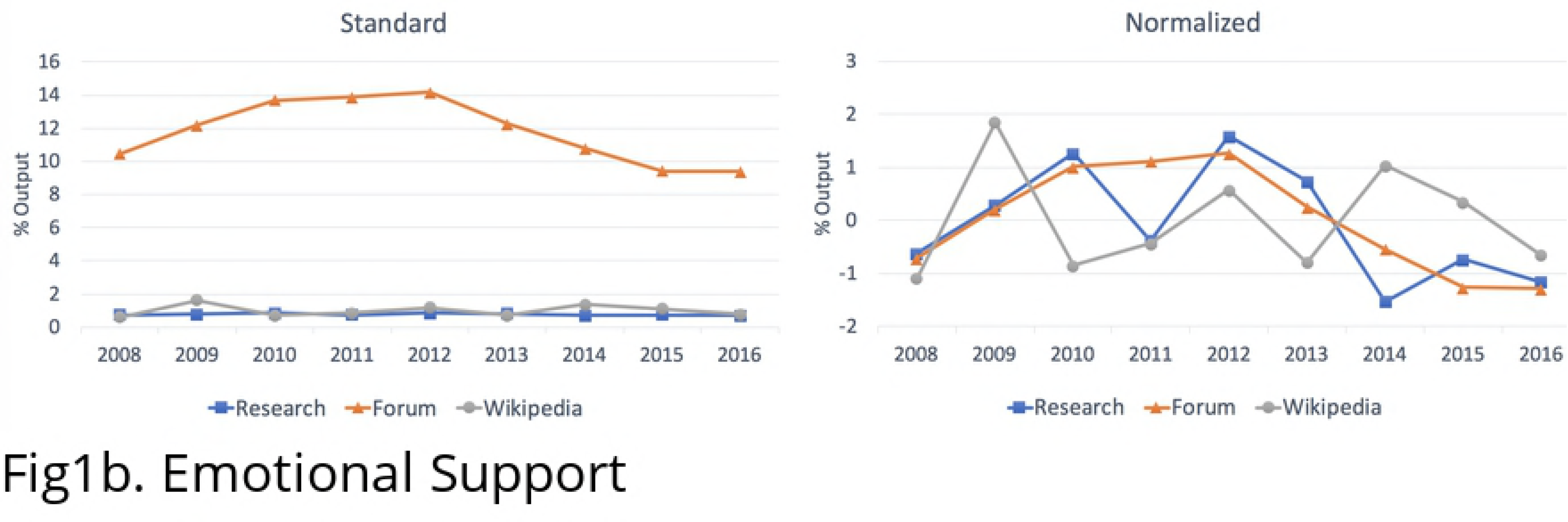
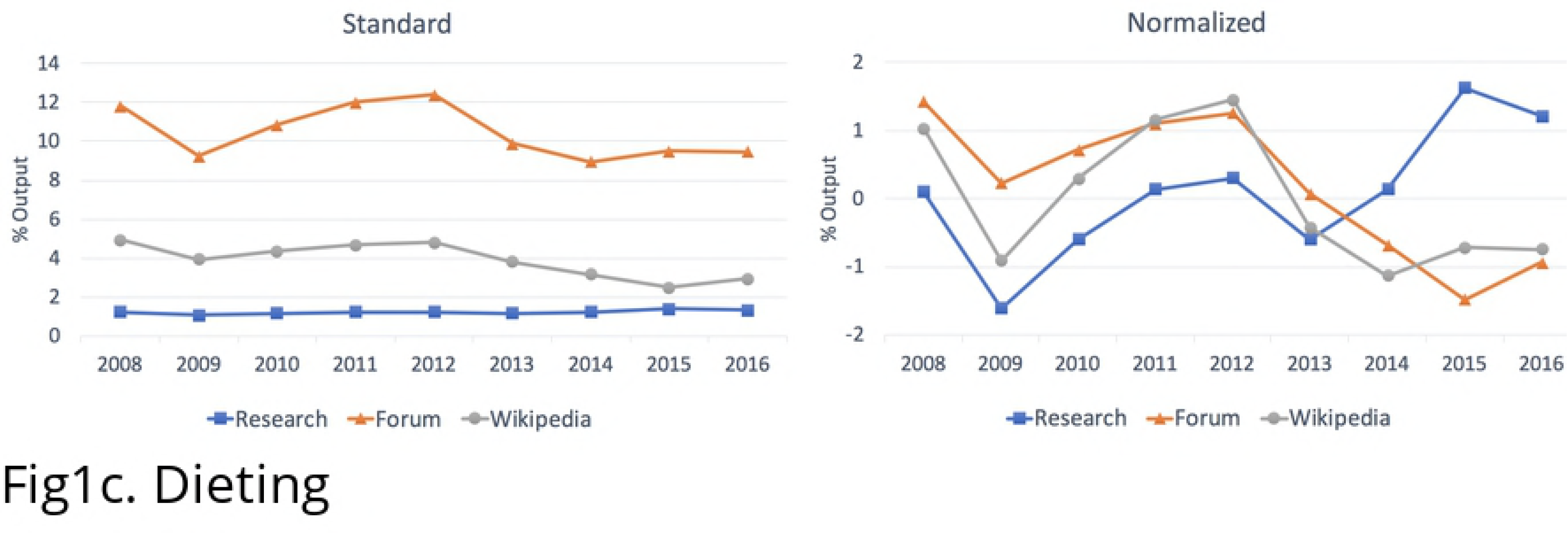
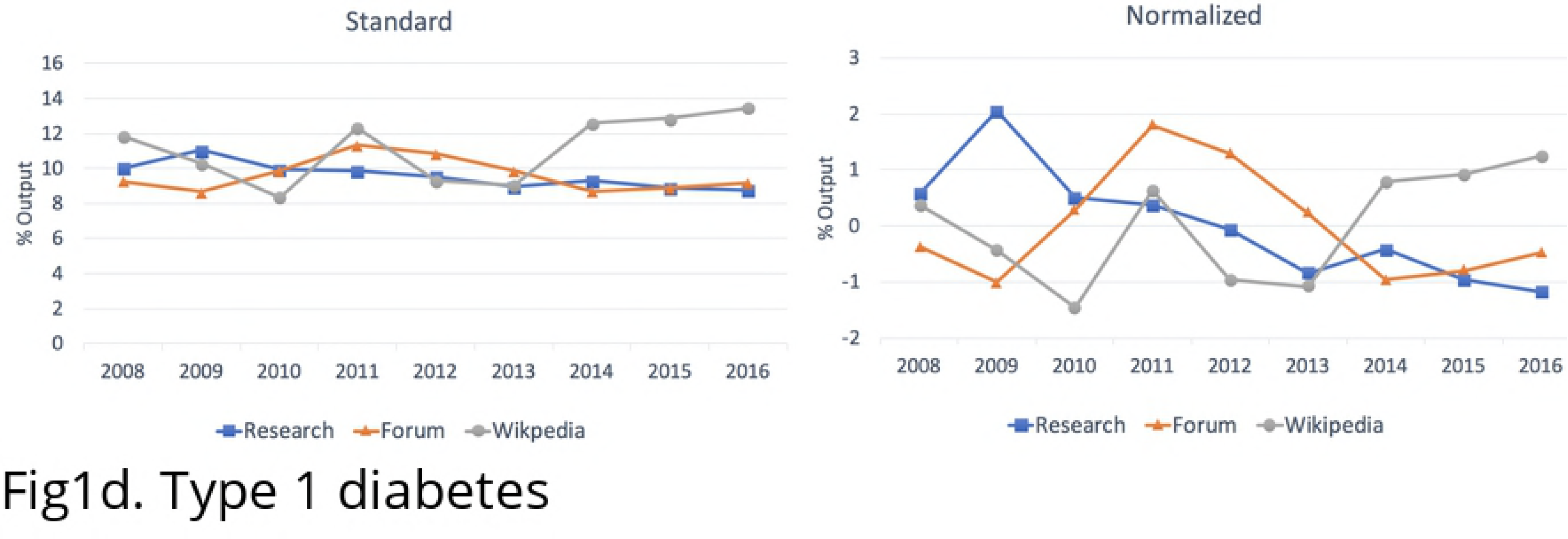
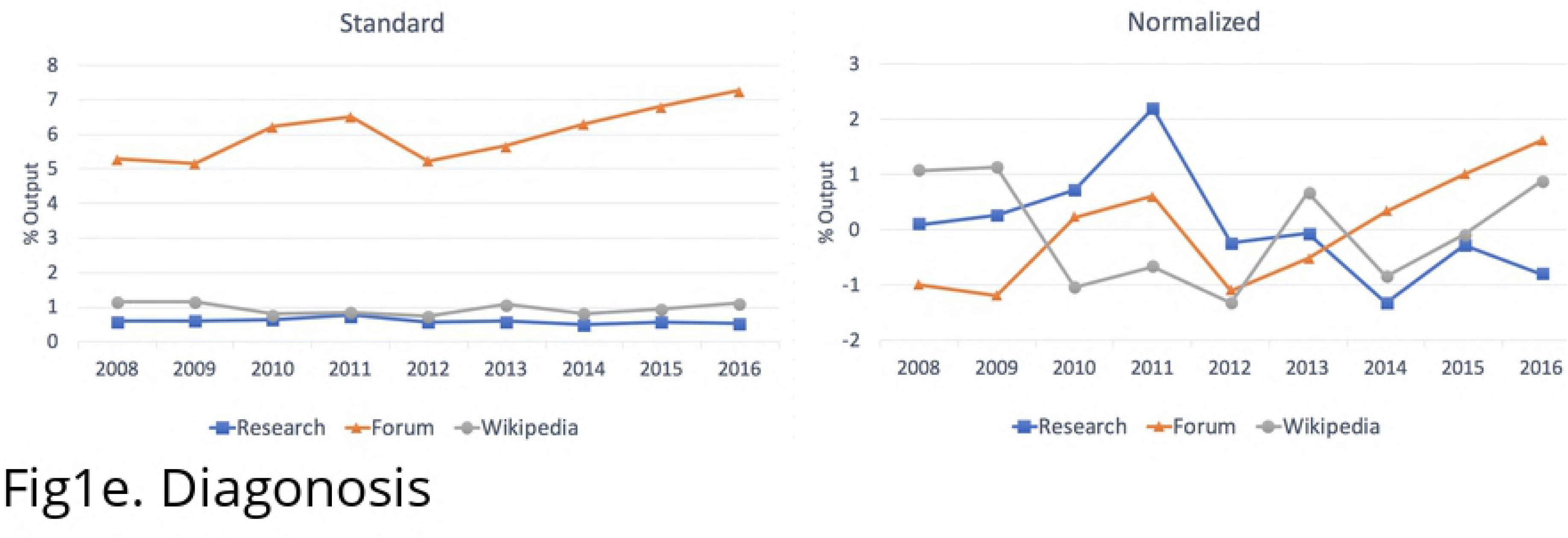
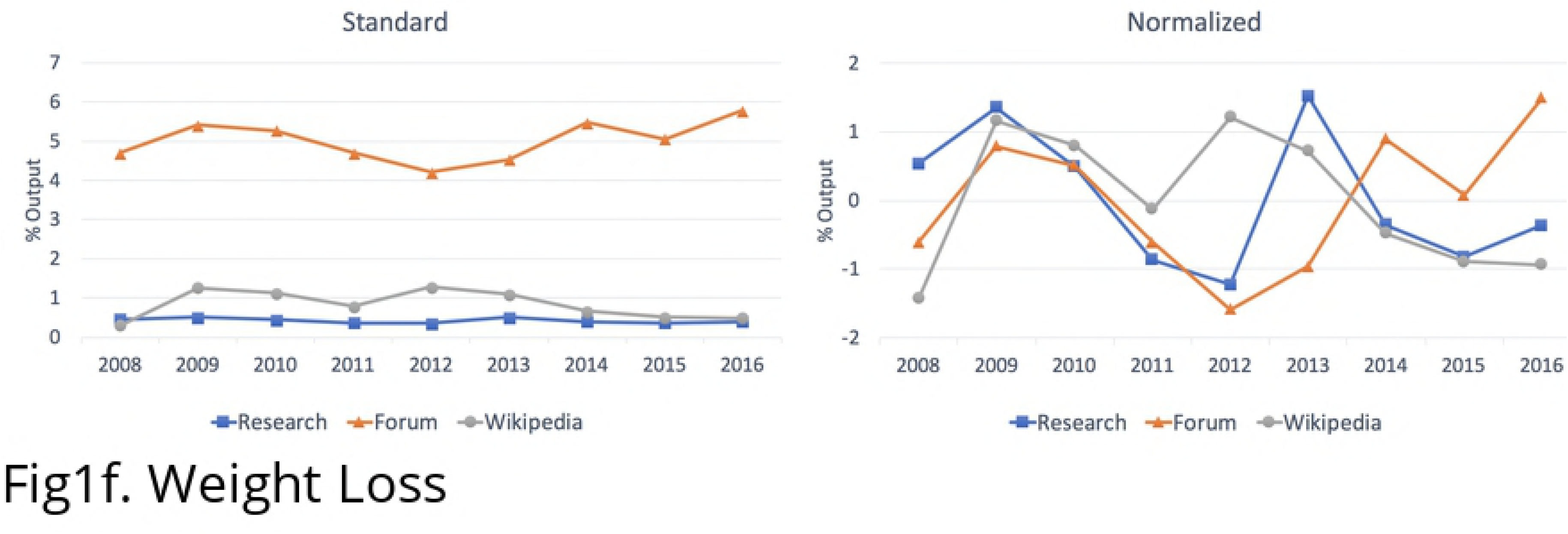
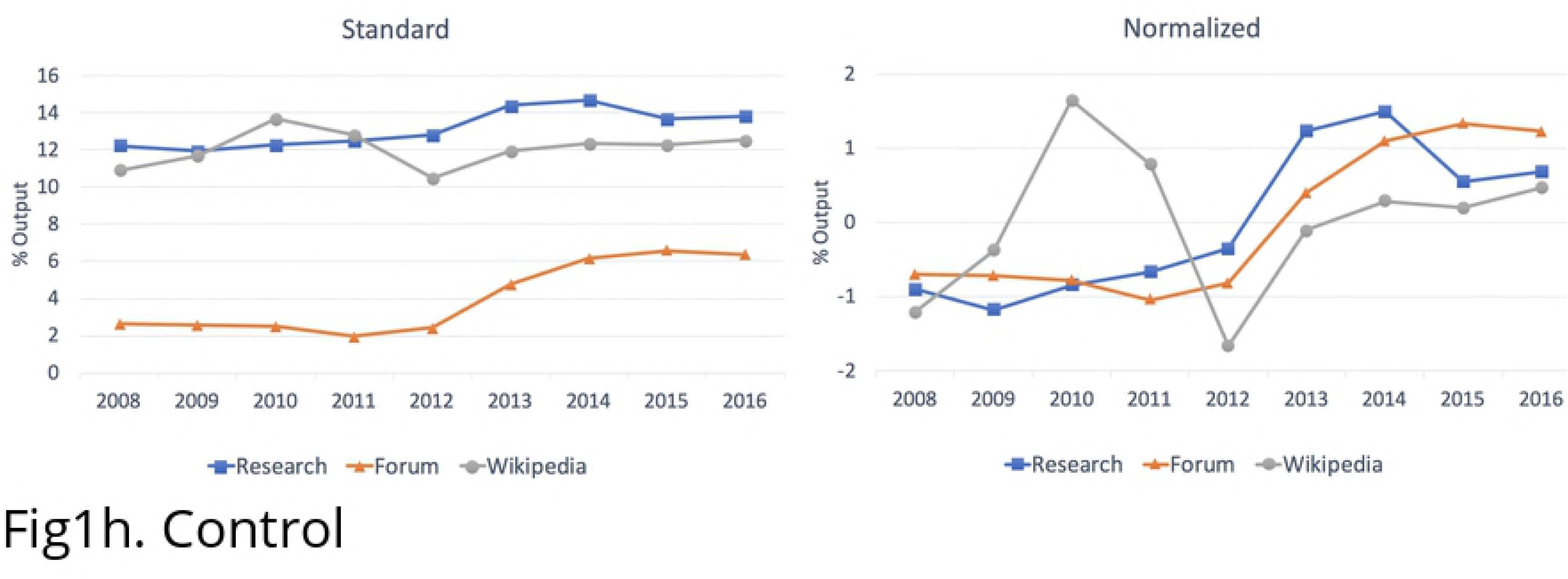
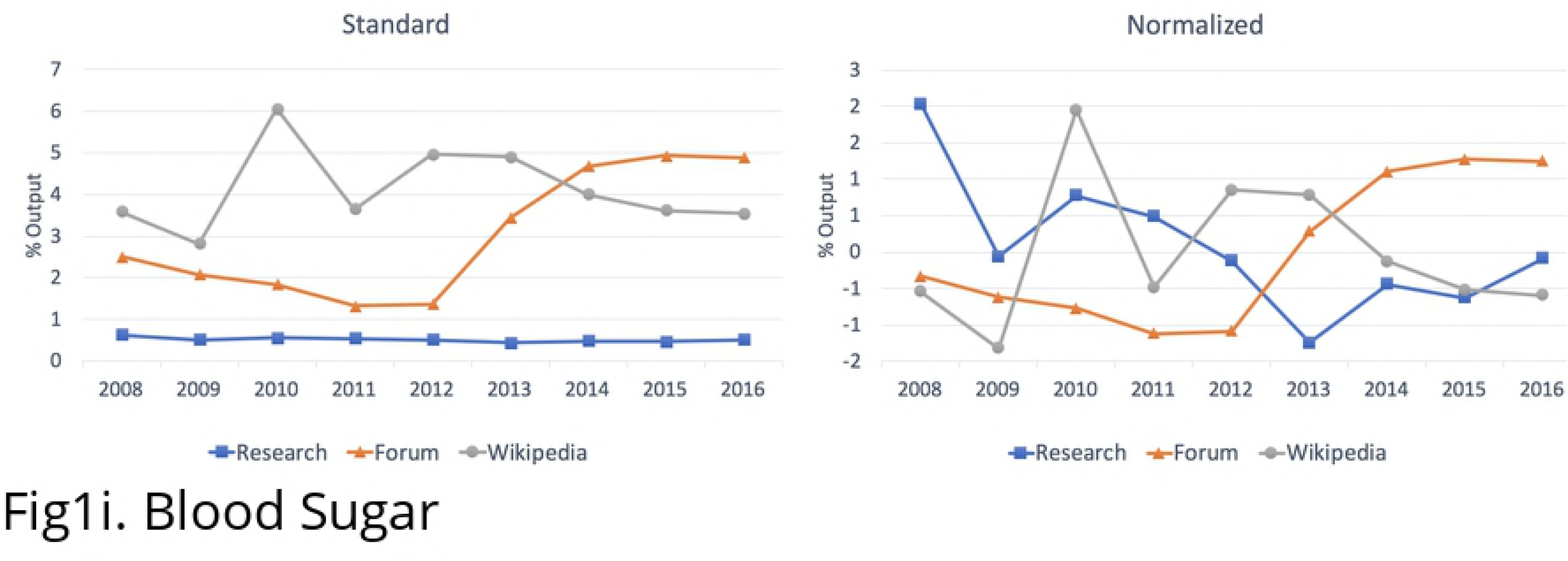
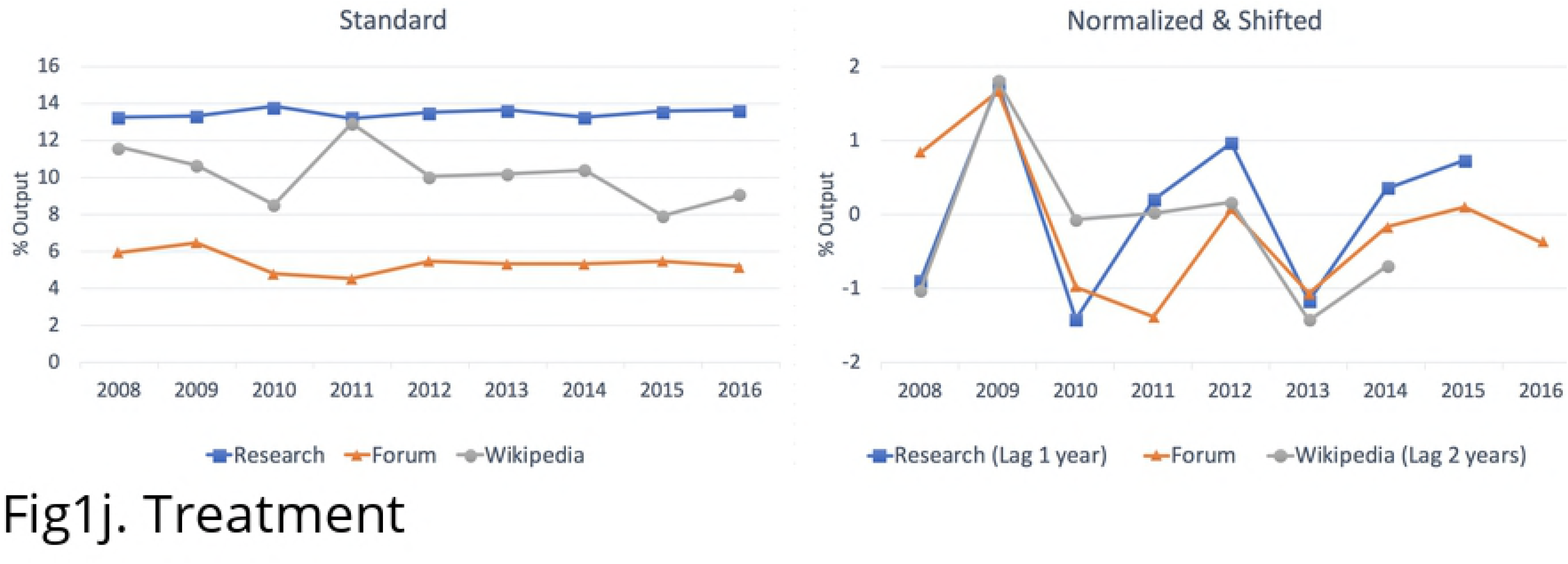
Percentage of research articles, Wikipedia articles, and forum discussions over years in the top ten forum topics.

## Discussion and conclusion

This case study explores the potential of using new sources, like public discussion forums, to better understand of the connection between the topics of interest to the public and those published in the research literature and edited on Wikipedia. The alignment in the rise and fall of relative interest over time in six of the top 10 topics (i.e., ‘emotional support’, ‘diet’, ‘type 1 diabetes’, ‘diagnosis’, ‘medication’, and ‘blood sugar’) is evidence that topic modeling approach used can be useful for comparing across platforms, even when these vary significantly in their function, users and language. In particular, it demonstrates the success of using the LLDA model as a strategy for classifying research and Wikipedia articles based on the language used by posts in a public community discussion forum. Moreover, the time-shifted alignment found in a couple of the series (i.e., type 2 diabetes and treatment) suggests that the method may be useful for detecting when topics gain popularity in one arena ahead of another. In these cases, rises and drops in interest by researchers happened both ahead of and after corresponding changes in relative interest in the discussion forum. The last two cases (weight loss/obesity and diabetes control) analyzed show that the method does not always find alignment between changes in relative interest in topics. These two cases reveal that the LLDA models are not mirroring each other but are reflecting varying degrees of interest in the topics across the three corpora.

Beyond comparisons to each other, several topics show a turning point in 2011 and 2012. The year 2012 specifically is an important date in the history of diabetes when death of 3.7 million diabetic patients was reported (WHO, 2012). The diabetes control trend increases across all platforms in 2012 and it continues increasing onwards (Figure 1h). A dramatic increase is also seen in diet trends in 2012 but the trends start dropping afterwards (Figure 1c). Similar trend is seen for emotional support where all the three trends peak at 2012 but starts decreasing afterwards (Figure 1b). This report received widespread attention and shifted the public conversation about diabetes (WHO, 2012; The Cleaner, 2014). Although we cannot make direct connections between the publishing of the report and the observed changes in topics of interest, this alignment is an encouraging sign that the LLDA approach applied to a discussion forum captures changes in public interests.

Using the LLDA method and assigning a more consistent set of topics to records in each platform eased and ensured the comparison task between a technical research corpus and a lay discussion corpus that has not been taken on in previous studies. In fact, no earlier study touched upon a lay discussion platform like a forum to detect community needs regarding a certain topic while such social platforms bear a great potential for detecting the most recent issues and concerns arising in a certain community. Previous research has instead focused on burden of diseases rates and global health trends as a way of measuring the public interest (Yegros, et al., 2018; Rafols & Yegros, 2018). In addition to research, project tools such as ‘*Searching for health*’ which was developed to show how Google searches for health information associate with actual incidences of diseases, does not shed lights on the alignment of online searches with the published research. Our proposed approach, piloted here, provides a more direct measure of a community’ interest, which could, in the future, be used in conjunction with the kind of public health data used in previous studies.

With regards to diabetes itself, and the analysis of diabetes related content, the results show that diabetes testing (especially the A1C test), diabetes control, and treatments are the most frequent topics in research articles indexed in Scopus and pages edited in Wikipedia while users of the discussion forum have a larger preoccupation with emotional support and motivation for patients and diabetes diet. Using the topics and the language found in the discussion forum, our topic model and time series show that the topics of published research and of the Wikipedia articles that are edited seem to largely correspond with one another. This shows that Wikipedia community follows scientific outputs more closely, something that is likely expected, given that Wikipedia editors are instructed to cite their sources using sources like peer-reviewed research in their work (Wikipedia, 2017) and that Wikipedia solicits scholars’ contributions (Corbyn, 2011). In this way, and to the extent that we studied it through the ten most popular topics, the results suggest that the content on Wikipedia is influenced and updated based on the conversations happening within the research community, more than the conversations happening between individuals affected by diabetes. The relationship between topics of interest on Wikipedia and in the research stand in contrast to the misalignment between them and the discussions taking place on the forum. This gap between public discussions and published research concurs with the results of previous studies showed no/little correlation between health issues based on disease burden rate and research published in academia (Rafols & Yegros, 2018; Yegros, et al., 2018; Evans, Shim, & Ioannidis, 2014).

However, although the topics do not thoroughly align between the forum and the research platforms, a time series analysis shows similar trends with similar peaks and downs from 2008-2016 for most topics including ‘emotional support’, ‘diet’, ‘type 1 diabetes’, ‘diagnosis’, ‘medication’, and ‘blood sugar’. In a nutshell, the three platforms examined, DiabeticConnect Forum, Wikipedia and Scopus, have different nature, so it is not expected to find a perfect alignment between their topics and their prevalence. It is logical that researchers are more interested in testing, control techniques and treatments than the general public as they are the experts responsible for the technical experiments and researching new methods with a wide access to laboratories and equipment. It is also logical for public to be mainly interested in their daily life issues such as emotions, mood, stress, support, food, etc. However, the misalignment found between research interest and public interest may be informative for the scientific community and health policy makers in diabetes to value some understudied areas that are of higher interest to public.

## Footnotes

1 http://www.searching-for-health.com

2 http://www.diabeticconnect.com/

3 https://en.wikipedia.org/w/api.php

4 http://mallet.cs.umass.edu/

